# Regional trait-threat interactions determine extinction risk in plants

**DOI:** 10.64898/2026.07.16.738776

**Authors:** Zoë Dennehy-Carr, Raphael LaFrance, Douglas E. Soltis, Pamela S. Soltis, Robert P. Guralnick

## Abstract

Global plant biodiversity and associated ecosystem services are being severely threatened by human activities and land-use change. This has led to an immense effort to assess the extinction risk of individual species and associated drivers to inform conservation actions and priorities. However, we lack a more general understanding of the processes that determine variation in vulnerability and resilience among plant taxa facing the same threats in the same or similar ecological conditions and geographic regions. This research helps identify these generalities and determines species and traits that are at disproportionate risk of extinction. Given Florida’s high levels of biodiversity and endemism, coupled with ongoing rapid and extensive human development, we use the state’s native flora as our model system.

By combining data on extinction risk, human-induced threats, and plant functional traits, our results show that species threatened with extinction cluster within trait space, with smaller herbaceous species at greater risk than larger woody species. We also find that all categories of threat considered in our study (Development and Infrastructure; Environmental Change and Modification; Natural Hazards and Climate Change; and Resource Extraction) increase extinction risk except for the ‘*Invasive Species, Pests, and Diseases’* threat category. Generalised linear mixed-effect models show differing responses of species to environmental change, habitat loss, and fragmentation, driven by differences in life-history strategies and dispersal capabilities and mechanisms. The varied responses of species facing the same threats uncovered here highlight that plant functional traits in the context of regional ecosystem processes can be used to predict species-specific responses to threats and land-use change, providing a valuable resource for conservation management and prioritisation.

**Article impact statement:** Plant functional traits can be used to estimate species-specific responses to human-induced threats and predict overall extinction risk.

## Introduction

Land plants (embryophytes) constitute ∼80% of global biomass and provide vital ecosystem services (Bar-On et al., 2018). Of the world’s ∼350,000 named vascular plant species, 30% have currently been assessed with an estimated 39 – 45% of those species threatened with extinction (Nic Lughadha et al., 2020; Govaerts et al., 2021; Antonelli et al., 2023), and a further unknown proportion is undescribed or have yet to be discovered (Brown et al., 2023; Ondo et al., 2024). Substantial losses of plant biodiversity and associated biomass will have profound effects on ecosystem service provision and functioning (Bachman et al., 2018), and consequently on the cultural and economic systems which depend upon them (Sato and Lindenmayer, 2017; Raven and Wackernagel, 2020; Bergstrom et al., 2021). In response, there has been significant investment in developing extensive infrastructure for assessing extinction risk, including the IUCN Red List (2026) and NatureServe (2026a), which provide species-level information on which species are imperilled and the threats that drive their extinction risk (Westwood et al., 2020; Carrero et al., 2022). Whilst these assessments inform much-needed conservation priorities and actions, they focus on documentation rather than understanding the underlying processes and characteristics determining species resilience and vulnerability across the plant tree of life.

Whether a species declines under a given threat depends on functional traits and life history strategies, and identifying these trait-based mechanisms is a key gap in understanding extinction risk (Chichorro et al., 2019, 2022; Green et al., 2022). For example, two co-occurring species may have different life histories and vary in functional trait space, resulting in different outcomes when facing identical threats. Without a more mechanistic account of why some species are vulnerable and others resilient to different threats, conservation prioritisation and actions remain reactive, protecting already imperilled taxa, rather than preventative, unable to anticipate which groups of currently secure species will decline following threat emergence (Álvare-Yépiz et al., 2019). This trait-based approach will also enable observed patterns to be generalised to unassessed and non-assessable taxa, floras, or future climates.

Closing this gap requires moving from documentation of patterns towards a more predictive, theory-driven understanding of plant vulnerability. Plant functional traits, including life-history strategy, growth form, and reproductive morphology, represent differing adaptations to environmental conditions and have long been used to explain how plants diversify, occupy new niches, and respond to environmental gradients (Taylor et al., 2023). For example, leaf succulence in *Agave* L. enables tolerance to extreme aridity and dominance in arid and semiarid North American ecosystems (Eguiarte et al., 2020), whilst carnivory in *Drosera* L. allows taxa to thrive in nutrient-poor, waterlogged soils (Williamson et al., 2025). Yet the same traits that confer an advantage under one set of environmental conditions can become disadvantageous under another, and extensive, rapid, human-driven disturbance is now reshaping the niche space available for plant lineages, in ways that may invert or alter the historical fitness of particular trait combinations (Vellend et al., 2017; Bruelheide et al., 2018; Feng et al., 2024). If variation in plant functional traits explains current patterns in biodiversity and diversification, the same variation should also explain species-level responses to human-induced disturbance. Several studies have begun to investigate the relationship between plant extinction risk and trait variation at coarse spatial scales (Pelletier et al., 2018; Gray, 2019; Humphreys et al., 2019; Ficken and Rooney, 2020). However, determining whether functional traits interact with specific threat types in driving extinction risk remains largely unaddressed, with most studies focusing on threats and risk separately rather than exploring their interacting effect (Butt and Gallagher, 2018; Ahler et al., 2023).

The flora of Florida is well-suited to addressing the potential interaction of traits and threats to extinction. The state occurs within the North American Coastal Plain biodiversity hotspot (Noss et al., 2015) with high levels of plant diversity and endemism, including ∼3000 native seed plants (WCVP, 2025). This diversity can be partially explained by Florida’s latitudinal gradient, which ranges from temperate to subtropical climates, encompassing a broad range of habitats such as cypress swamps, longleaf pine woodlands, and mangrove forests across three EPA level III Ecoregions (Southeastern Plains, Southern Coastal Plain, and Southern Florida Coastal Plain; Griffith et al., 2013). Changing sea levels throughout the Pleistocene also led to repeated periods of isolation and contact, driving lineage radiations and endemism (Naranjo et al., 2022; Nevado et al., 2024). These environmental and climatic factors have also led to the evolution of a wide range of plant functional traits and life-history strategies.

Florida’s unique and diverse flora is undergoing rapid, well-documented species loss, with much of the state’s biodiversity and ecosystems threatened with extinction, including 457 seed plant species (NatureServe, 2026). This threat is primarily due to rapid statewide development and associated land-use change, following continued increases in human populations, which rose 16.34 % alone between 2015 and 2025 (US Census Bureau, 2026). This rapid increase in population has resulted in extensive habitat degradation, fragmentation, and loss, including threats to important habitats such as scrub oak woodlands and prairie grasslands, upon which many endemic species depend (Lucash et al., 2022). The flora is also highly threatened by invasive and naturalised species (Stohlgren et al., 2006; Lieurance et al., 2023) and the increasing frequency of stochastic natural hazards such as droughts and hurricanes (Reece et al., 2013; Lucash et al., 2022; Gonçalves et al., 2024). Consequently, Florida is a region in which multiple, distinguishable anthropogenic and environmental threats are present across a large spatial scale, affecting a taxonomically and functionally diverse flora.

Using the Florida flora as a model system, we test whether plant functional traits, threats, and their interactions explain variation in species extinction risk. We predict traits associated with slow life-history strategies will be associated with elevated risk, particularly in the presence of threats which initiate land clearance and modification, whilst traits associated with broad environmental tolerances and fast life histories will be neutral or at reduced risk. By moving the analysis from a traits-and-risk to a trait-threat-risk interaction framework, we shift the framing of plant vulnerability from a descriptive question (which species are at risk?) toward a mechanistic one (which traits, under which threats, predict risk?). Our framework can be applied to other floras of varying geographical extents and taxonomic groups for which comparable assessment and trait data exist.

## Materials and Methods

### Florida seed plant checklist

To generate our species list of Florida plants, we followed the method of Carruthers et al. (2026 [preprint]) for taxonomic reconciliation with names standardised according to the World Checklist of Vascular Plants (WCVP; Govaerts et al., 2021) and intraspecific taxa represented at the species level. Native taxon distributions from WCVP were used to construct a list of all native seed plant species known to occur in Florida. We used the most up-to-date WCVP v.14 taxonomic backbone and species distributions (Govaerts et al., 2025; https://doi.org/10.34885/b8fr-km05).

### Trait Data

We collated data for 17 functional traits from 14 sources (Supplementary Tables 1 and 2) to determine how key floristic traits affect extinction risk probability in angiosperms and gymnosperms. Data from the *Flora of North America* (FNA) were acquired using rule-based parsing for taxon accounts published online and provisional or unpublished accounts for 14 families (Supplementary Table 3). Trait databases BIEN, GIFT, and TRY were primarily used for categorical data that could not be readily acquired from FNA (Supplementary Table 1). Data from BIEN (Maitner et al., 2017) and GIFT (Denelle et al., 2023) were downloaded using their respective R packages, whilst the online TRY data portal was used to request data (Kattge et al., 2020). Trait data from other sources were manually extracted. For data integration, a single value was assigned to each species. Given its expert curation and description, data obtained from FNA were selected over all other data sources. Mean seed mass values were obtained from the Seed Information Database. For all other continuous data, the maximum value for each species was selected, unless it was identified as an outlier value. In such cases, trait values were manually verified using other data sources to reach a consensus. A similar process occurred for categorical data, with differences reconciled using a third source prior to data integration. All data sources are listed in Supplementary Table 2. Following data integration and cleaning, coverage of each trait was calculated with poorly represented traits excluded from our analyses to reduce the effect of missing data.

### Conservation Status and Threat Data

Data on conservation status and threats were acquired from NatureServe using rule-based parsing. NatureServe conducts assessments on species native to the United States and North America, providing data on conservation risk and biodiversity threats. Species are assigned global status ranks from G1 (critically imperilled) to G5 (secure), and GX (presumed extinct) or GH (possibly extinct) (Supplementary Table 4). For this study, species assigned G1-G3 are considered ‘at risk’, and species ranked G4 or G5 are considered ‘secure’ and used to assess extinction risk. NatureServe uses the IUCN-CMP Threat classification scheme v.3.2 (IUCN, 2012), as a result, we follow the definitions of this classification scheme rather than the most recent version (v.4.0; Salafsky et al., 2025) to ensure consistency. The IUCN threat classification scheme (v.3.2) includes a total of 11 Level 1 threat categories, which we grouped into five broader groups based on the impact and mechanism of each threat (Table 1; Supplementary Table 5). Species were assigned threat categories from free-text descriptions found under the fields of ‘NatureServe Status’ and ‘NatureServe Global Conservation Status Factors’.

**Table 1.** Threat classifications used in this study and their corresponding IUCN-CMP level one categories (IUCN, 2012).

| **Threat** | **IUCN-CMP Level One Categories** |
| --- | --- |
| **Development and Infrastructure** | 1. Residential & Commercial Development<br>4. Transportation & Service Corridors |
| **Environmental Change and Modification** | 6. Human Intrusions & Disturbance<br>7. Natural System Modifications<br>9. Pollution |
| **Invasive Species, Pests, and Diseases** | 8. Invasive and Other Problematic Species, Genes and Diseases |
| **Natural Hazards and Climate Change** | 10. Geological Events<br>11. Climate Change and Severe Weather |
| **Resource Extraction** | 2. Agriculture & Aquaculture<br>3. Energy Production & Mining<br>5. Biological Resource Use |

### Principal Component Analysis (PCA)

Given possible trait collinearity, we ran principal component analyses (PCA) to produce reduced uncorrelated trait axes and determined how scores of species on these axes affect plant extinction risk and the impact of different threats. To assess the impact of removing seed mass on trait variability, we ran two PCAs, first using matrix A which includes two categorical (growth form and life cycle) and five continuous traits (fruit length, leaf length, leaf width, plant height, and seed mass) and then using matrix B which includes the same two categorical traits and four continuous traits with seed mass removed. All species without a corresponding NatureServe assessment or that did not have trait data recorded for all continuous traits selected were excluded from our analyses. Values for all traits were natural log-transformed, and the PCA was fitted using the function *PCA* in the R package *FactoMineR* (Lê et al., 2008) and visualised using the *factoextra* R package (Kassambara and Mundt, 2026). PC scores on these axes were used downstream in models described below.

### Generalised Linear Mixed-Effect Models

We used generalised linear mixed-effect models with a binomial distribution to test how functional traits and threats relate to species extinction risk. Extinction status was binarised as the response variable (secure = 0, at risk = 1). Continuous predictors (PC1 and PC2 scores) were standardised using the *scale* function in R. Growth habit (herbaceous or woody) and life cycle (annual, biennial, perennial, or variable) were included as categorical fixed effects in all models. Plant family (following APG IV) was included as a random intercept to account for phylogenetic relatedness and clustering. Models were fitted using the *glmer* function in the *lme4* R package (Bates et al., 2015), using the bobyqa optimiser to aid convergence.

We compared six candidate models that represent distinct hypotheses about how single and multiple threats, along with their interactions with plant functional traits may shape extinction risk. Our first model included only traits and tested whether functional traits alone predict extinction risk. The remaining five models added the threats examined here and traits in different combinations with increasing complexity. Model two included the cumulative number of threats as a continuous predictor, testing whether total threat burden scales linearly with risk. Model three included the cumulative number of threats as a factor, allowing for non-linear effects of threat accumulation. Model four included a single binary term for the presence of any threat, testing whether the presence rather than the number of threats matters. Model five included each of the five threat categories (Table 1) as separate binary predictors (absence = 0, presence = 1), testing whether specific threats differ in their effects. Model six extended model five by adding interaction terms between PC1 and PC2 and the two threat categories most likely to interact with plant traits (i.e., Development and Infrastructure and Environmental Change and Modification).

The best model was selected using AIC and AIC weights and likelihood tests. We assessed multicollinearity in the best-fitting model using *check_collinearity* in the *performance* R package (Lüdecke et al., 2021) and by calculating the variance inflation factor (VIF) using the *car* R package (Fox and Weisberg, 2026). We quantified the variance explained by fixed effects using the *r2_nakagawa* function from the same package. Predicted probabilities of being at risk were calculated using *ggpredict* in the *ggeffects* R package (Lüdecke et al., 2018), and odds ratios for all threat categories were also calculated.

## Results

### Conservation Status and Threat Data

Approximately 75% of the 2,989 plant species native to Florida are considered secure (G4 and G5), ∼15% at risk (G1-G3), and three are either presumed extinct (GX) or possibly extinct (GH) by NatureServe (Table 2). Less than 10% of species native to Florida have not yet been assessed. Among the 2,702 extant species assessed by NatureServe, 26.76% are affected by at least one threat (Table 3). This includes 17.68% of secure species and 71.33% of at-risk species. The most prevalent threat type across all groups is ‘*Development and Infrastructure*’ (20.28% of all species, 13.41% of those considered secure, and 54.05% of those at risk), followed by ‘*Environmental Change and Modification*’ (16.51 % of all species) and ‘*Invasives, Pests, and Diseases*’ (13.95% of all species).

**Table 2.** The proportion of species displayed as a percentage by NatureServe category for all species recorded to be native to Florida (not assessed includes both NatureServe categories GNR and GU, as well as species where we could not recover any listing).

| **NatureServe Category** | **Proportion of species (%)** | **Conservation Status** | **Proportion of species (%)** |
| --- | --- | --- | --- |
| **G1** | 2.54% (76) | **At-Risk** | 15.29% (457) |
| **G2** | 3.71% (111) |  |  |
| **G3** | 9.03% (270) |  |  |
| **G4** | 23.75% (710) | **Secure** | 75.11% (2245) |
| **G5** | 51.36% (1535) |  |  |
| **GH/GX** | 0.09% (3) | **GH/GX** | 0.09% (3) |
| **Not Assessed** | 9.50% (284) | **Not Assessed** | 9.50% (284) |

**Table 3.**
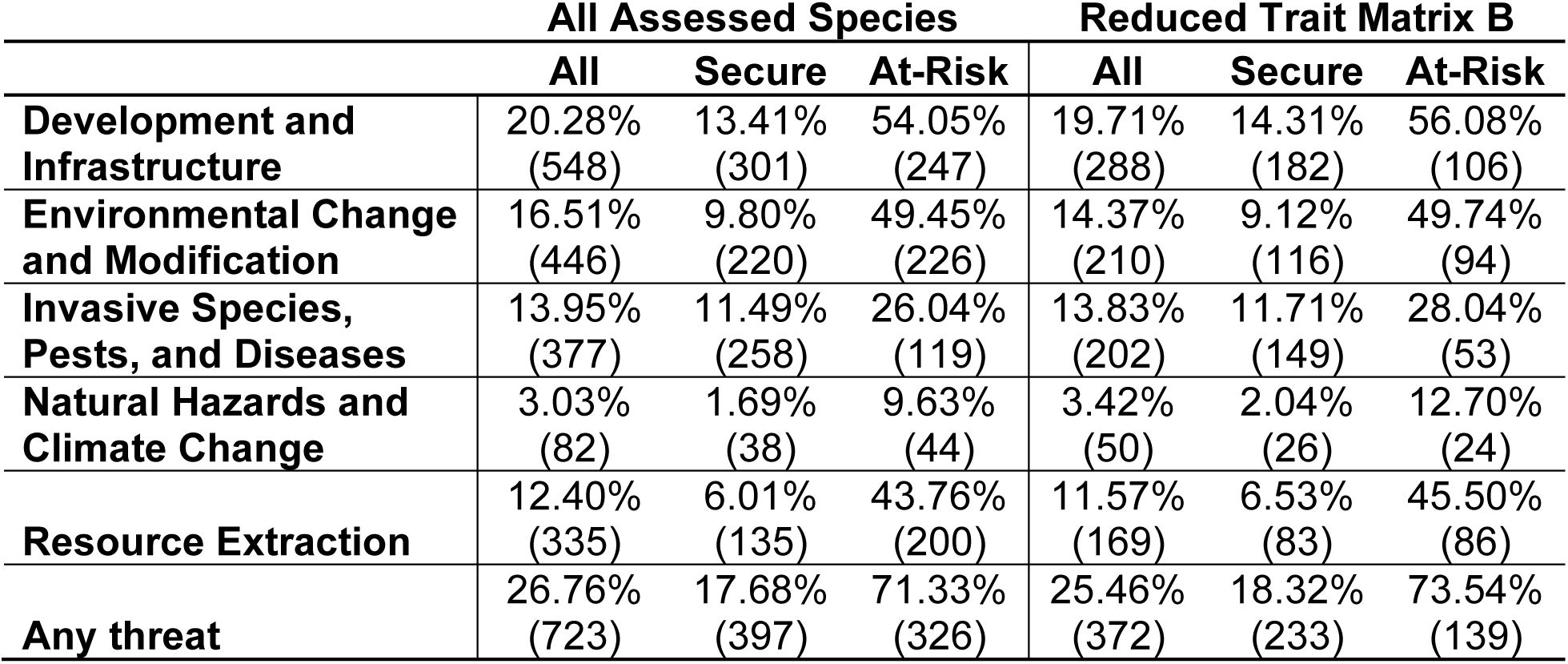
The number of species recorded to be impacted by each threat category for all species included in our trait matrix and all species assessed by NatureServe in Florida, USA.

### Trait Coverage

Due to high levels of missing and/or erroneous trait data, the number of traits was reduced for our analyses (Table 4). Out of the 2,702 species with NatureServe assessments, 691 species had data for all traits in matrix A (fruit length (mm), growth form, leaf length (cm), leaf width (cm), life cycle, plant height (cm), and seed mass (g)) compared to 1,461 species in matrix B (fruit length (mm), growth form, leaf length (cm), leaf width (cm), life cycle, and plant height (cm)). We selected matrix B for our analysis, with eight species classed as at risk included within matrix A, but we ran comparative PCA analyses to confirm the effect of removing seed mass. Within matrix B, all continuous traits had >77% coverage. None of the gymnosperm species that are native to Florida have data for all traits and therefore, are removed from our analyses, which will subsequently focus on angiosperm species only.

**Table 4.** Representation of plant trait data used in this study for all native species and species with NatureServe assessments of G1-G5, broken down by conservation status. The proportion of species in matrix A (growth habit, life cycle, fruit length, leaf length, leaf width, plant height, and seed mass) and matrix B (growth habit, life cycle, fruit length, leaf length, leaf width, and plant height) are included for comparative purposes.

| Trait | Proportion of all species (%) | Proportion of assessed species |  |  |
| --- | --- | --- | --- | --- |
|  |  | All species (%) | Secure species (%) | At-risk species (%) |
| **Life Cycle** | 99.93% (2987) | 99.96% (2701) | 99.96% (2244) | 100.00% (457) |
| **Growth Habit** | 99.63% (2978) | 99.81% (2697) | 99.82% (2241) | 99.78% (456) |
| **Fruit Length (mm)** | 76.55% (2288) | 77.57% (2096) | 78.80% (1769) | 71.55% (327) |
| **Leaf Length (cm)** | 83.61% (2499) | 84.64% (2287) | 85.70% (1924) | 79.43% (363) |
| **Leaf Width (cm)** | 79.59% (2379) | 81.20% (2194) | 82.76% (1858) | 73.52% (336) |
| **Plant Height (cm)** | 84.18% (2516) | 85.20% (2302) | 87.66% (1968) | 73.09% (334) |
| **Seed Mass (mg)** | 39.98% (1195) | 41.78% (1129) | 49.27% (1106) | 5.03% (23) |
| **Matrix A** | 26.73% (799) | 25.87% (699) | 30.78% (691) | 1.75% (8) |
| **Matrix B** | 52.86% (1580) | 54.07% (1461) | 56.66% (1272) | 41.36% (189) |

### Principal Component Analysis (PCA)

Variation in our trait dataset is predominantly explained by principal component one (PC1), which accounts for 48.9% of the variation (Figures 1,2). All four continuous traits from matrix B are positively correlated with PC1 (Table 5), which represents plant size: species with higher PC1 values are taller and have larger leaves (leaf length and width) and longer fruits than those with lower PC1 values. Principal component two (PC2) represents 24.3% of the variation and is explained by differences in leaf and fruit lengths (63.3% and 33.3% contribution, respectively). Leaf length is strongly positively correlated with PC2, whilst fruit length shows a moderate negative correlation, indicating species with high PC2 values have longer leaves and shorter fruits.

**Figure 1.**
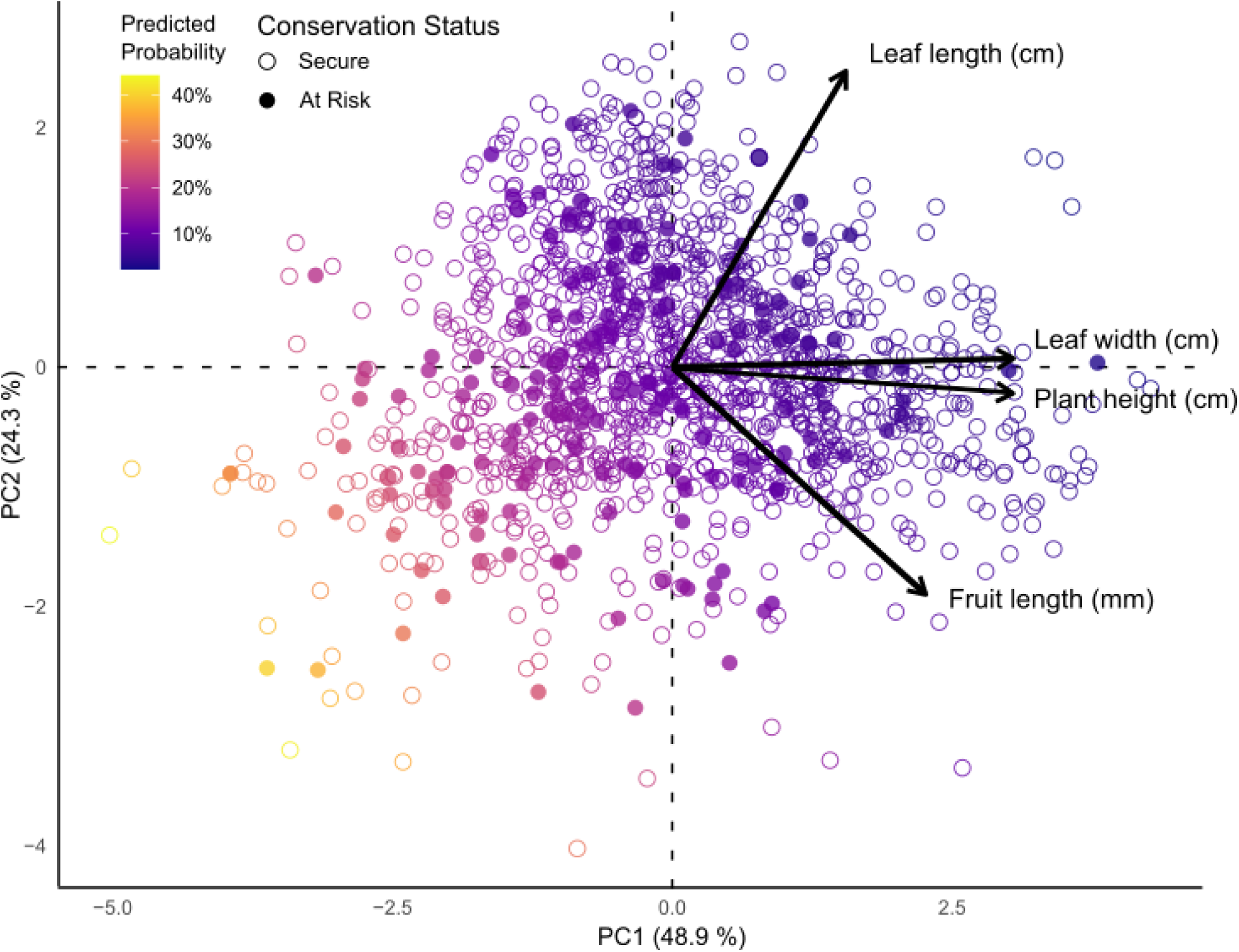
Principal component analysis biplot of trait variation for 1,461 plant species native to Florida. Continuous trait values are logged: leaf length (cm), leaf width (cm), plant height (cm), and fruit length (mm). Each circle represents one species, with filled circles representing at-risk species (G1-G3) and open circles representing secure species (G4-G5) according to their corresponding NatureServe assessments. The colour gradient indicates the predicted probability of extinction risk.

**Figure 2.**
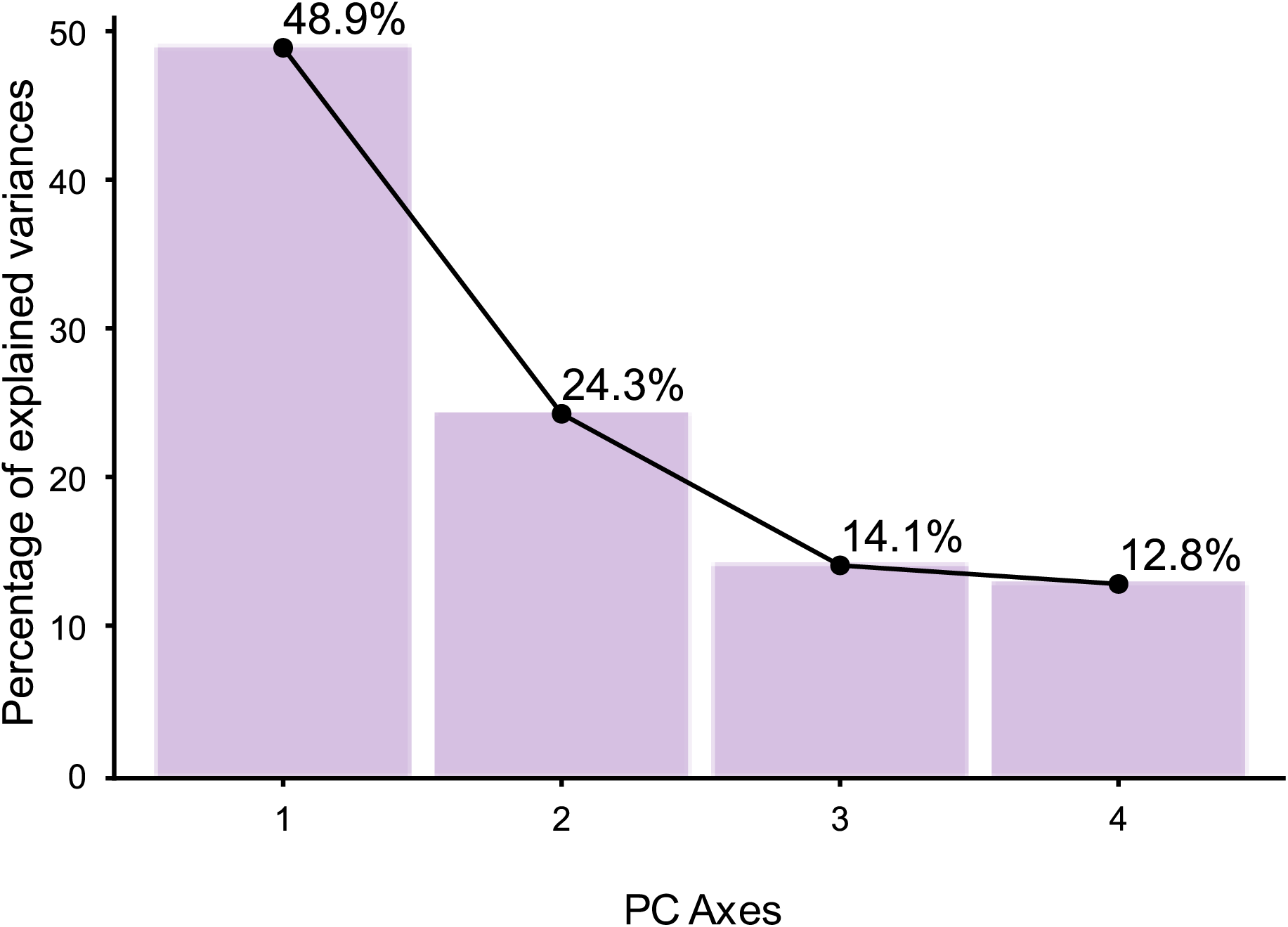
Principal component analysis eigenvalue scree plot of trait variation for 1,461 species native to Florida. Trait loadings for each PC axis are presented in Table 5 below.

**Table 5.**
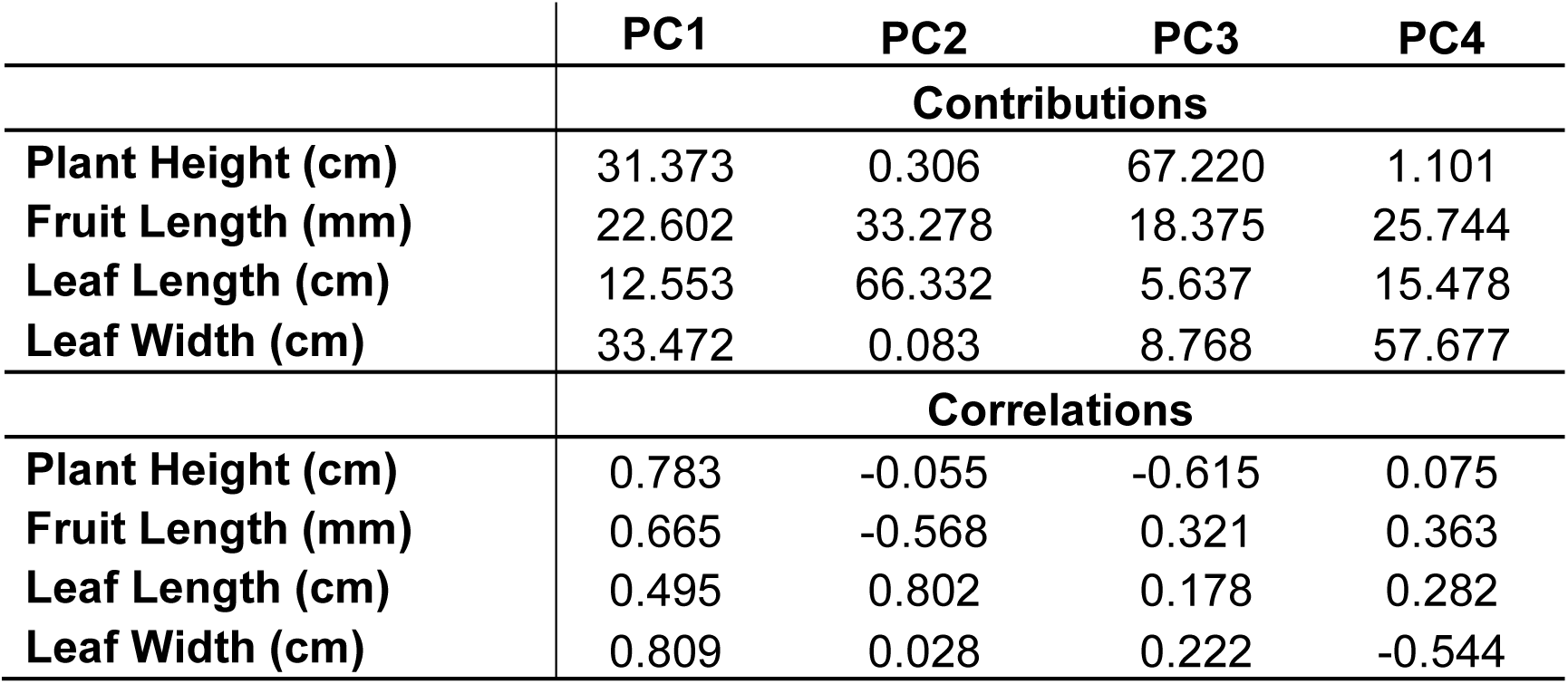
The contributions and correlations of each continuous trait with each principal component for 1,461 angiosperm species native to Florida.

|  | **PC1** | **PC2** | **PC3** | **PC4** |
| --- | --- | --- | --- | --- |
|  | **Contributions** |  |  |  |
| **Plant Height (cm)** | 31.373 | 0.306 | 67.220 | 1.101 |
| **Fruit Length (mm)** | 22.602 | 33.278 | 18.375 | 25.744 |
| **Leaf Length (cm)** | 12.553 | 66.332 | 5.637 | 15.478 |
| **Leaf Width (cm)** | 33.472 | 0.083 | 8.768 | 57.677 |
|  | **Correlations** |  |  |  |
| **Plant Height (cm)** | 0.783 | -0.055 | -0.615 | 0.075 |
| **Fruit Length (mm)** | 0.665 | -0.568 | 0.321 | 0.363 |
| **Leaf Length (cm)** | 0.495 | 0.802 | 0.178 | 0.282 |
| **Leaf Width (cm)** | 0.809 | 0.028 | 0.222 | -0.544 |

To determine the impact of not including seed traits in our analyses, we ran a PCA analysis using matrix A, which included seed mass as an additional trait (Supplementary Figures 1, 2; Supplementary Table 6). This analysis included 691 species that have been assessed by NatureServe. The relationships and variance explained between both analyses are similar, with PC1 and PC2 accounting for 48.6% and 21.2% of the variation, respectively, and both principal components represented by the same traits (Table 5, Supplementary Table 6). Seed mass is positively correlated with PC1 and negatively correlated with PC2, aligning between fruit length and plant height (Supplementary Figure 1). Due to the low coverage of seed mass data, matrix A only includes eight species considered to be at risk by NatureServe. Considering this limitation and the similar patterns of trait variation recovered between these PCA analyses, we use PC1 and PC2 values from our PCA analysis constructed using matrix B for our generalised linear mixed-effect models to adequately test our hypotheses.

### Generalised Linear Mixed-Effect Models

We used a general linear modelling framework to ask if traits and threats interact to determine if a species is more likely to be considered at risk. Model one tests whether the functional traits included can predict plant extinction risk regardless of the presence or absence of different threats. Models two to six tested the effect of threats on extinction risk with increasing complexity. The best-fitting model was selected using AIC values and likelihood tests. Model six was identified as the best-fitted model, with the lowest AIC value (Supplementary Table 7) and has an improved fit over the non-interaction model five confirmed using a likelihood ratio test. Model six tests whether the presence or absence of different threats can predict plant extinction risk. Additionally, this model includes interactions between traits and threats on the effect of plant extinction risk to determine if specific plant traits make species either more or less vulnerable to specific threats. Non-significant threat interaction terms were removed to improve model fit. The structure of model six is presented below:

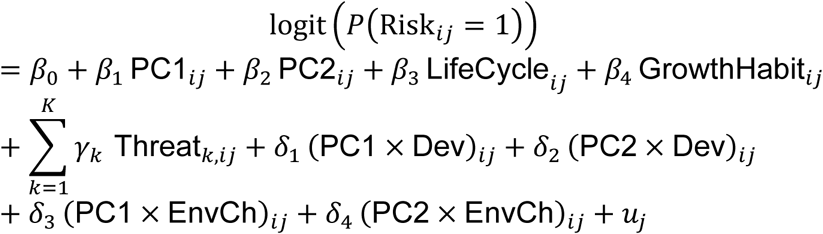

with 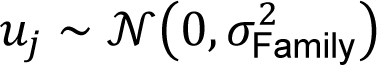. Each 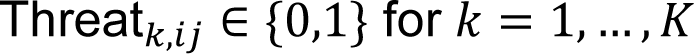 threat categories. Dev denotes Development and Infrastructure. EnvCh denotes Environmental Change and Modification.

Model one shows a significant increase in the probability of extinction risk with decreasing PC1 and PC2 values (Figure 3; Table 6). As PC1 and PC2 values decrease from 0 to -2 and -4, the predicted probability of extinction risk increases from 5% to 21% and 60%, respectively with smaller, herbaceous species exhibiting an increased probability of extinction compared to larger, woody species.

**Figure 3.**
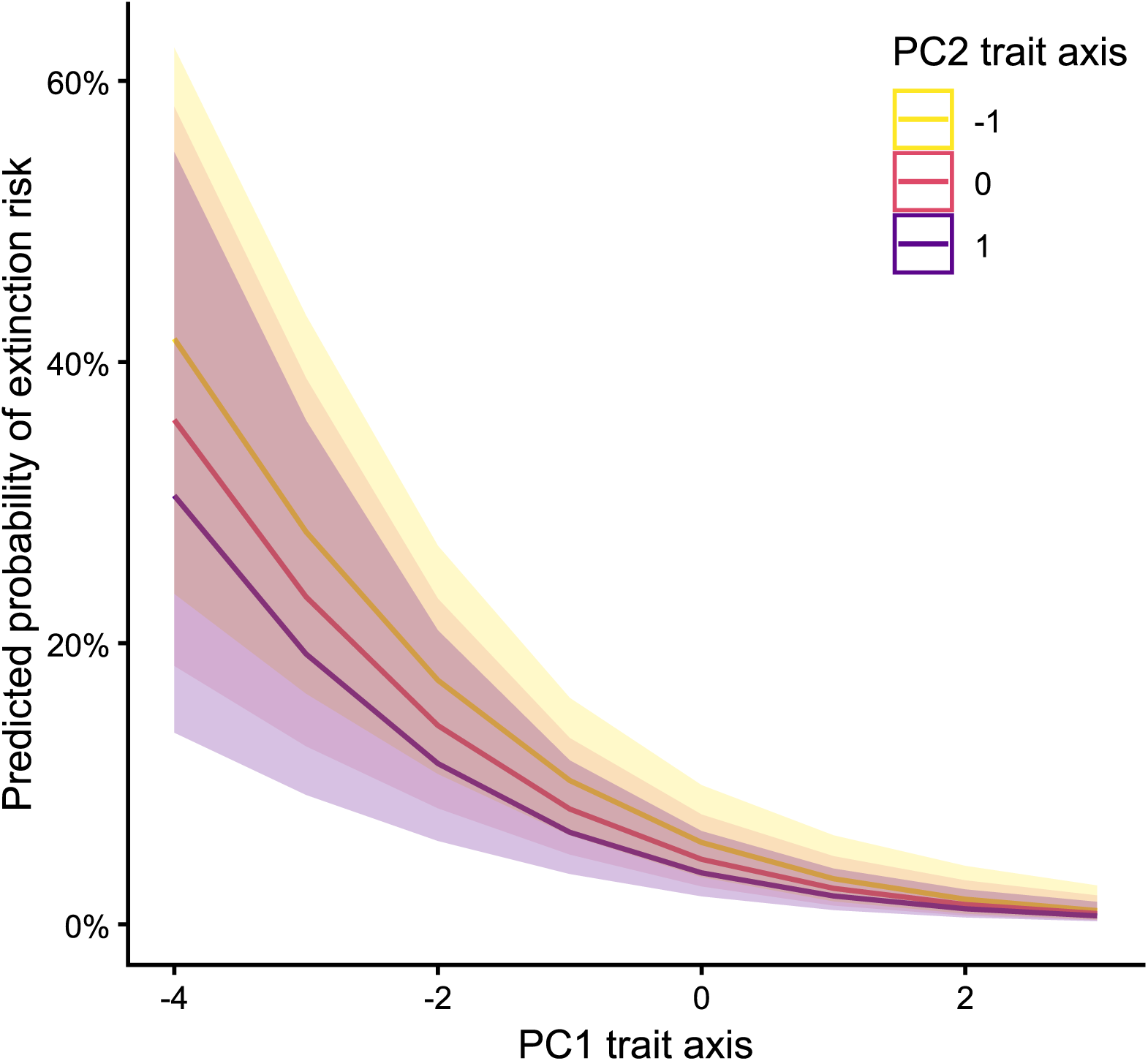
Generalised linear mixed-effect models showing the effect of PC1 and PC2 values on the predicted probability of extinction risk.

**Table 6.**
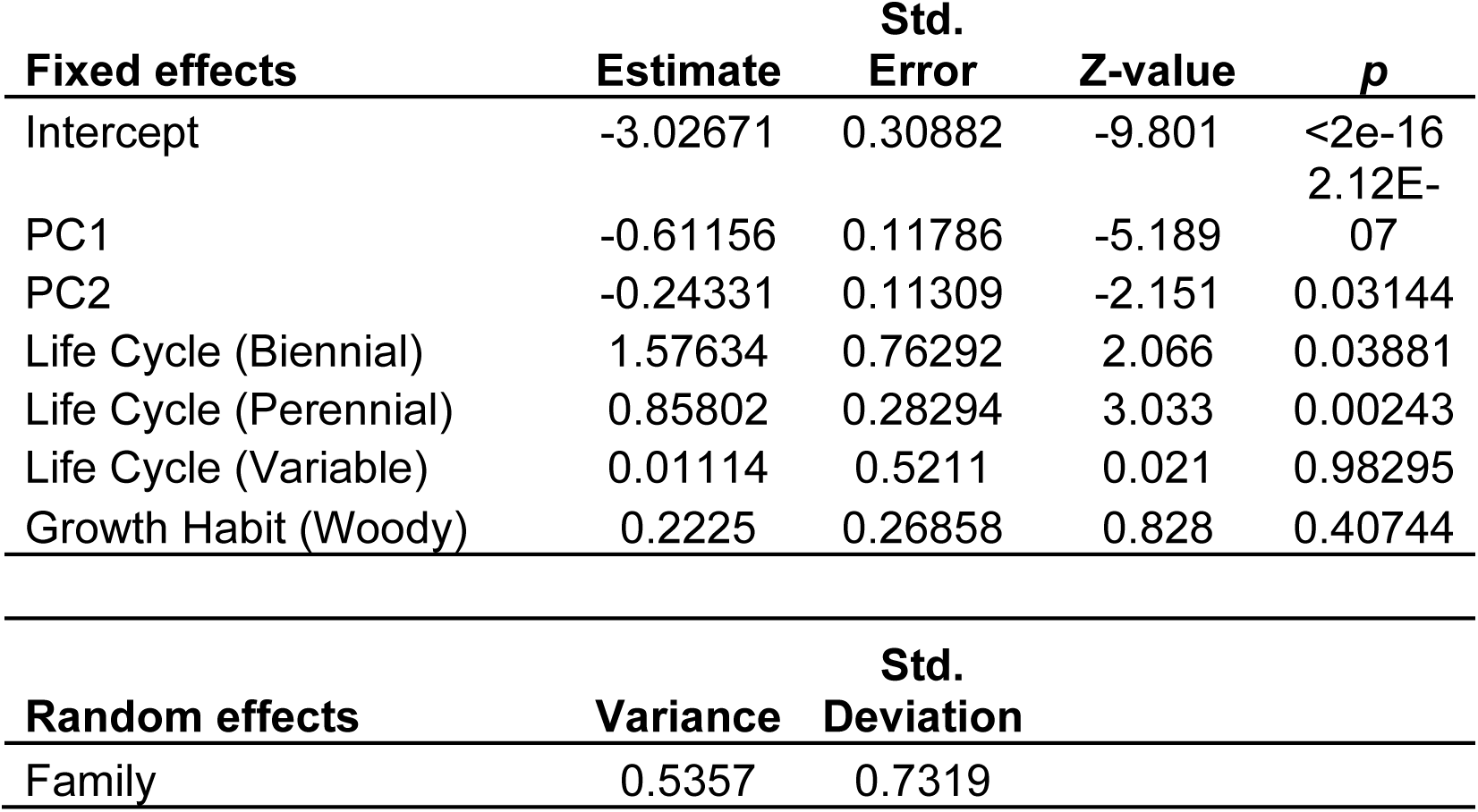
Summary results of generalised linear mixed-effect model predicting the effect of PC values on the probability of extinction risk.

Model six shows all threat types significantly predict extinction risk, with ‘*Resource Extraction’* (odds ratio (OR) = 5.29; 95% CI: 3.04-9.21) and ‘*Environmental Change and Modification’* (OR = 4.23; 95% CI: 2.37-7.55) having the strongest effects (Figure 4, Table 7). The presence of *‘Invasive Species, Pests, and Diseases*’ was the only threat to reduce the probability of extinction risk (OR = 0.47; 95% CI: 0.26-0.83), whilst the presence of all other threats increased the odds of extinction risk. The results of all models constructed are presented in Supplementary Table 8. For both models one and six, VIF values were <1.6 (Supplementary Table 9).

**Figure 4.**
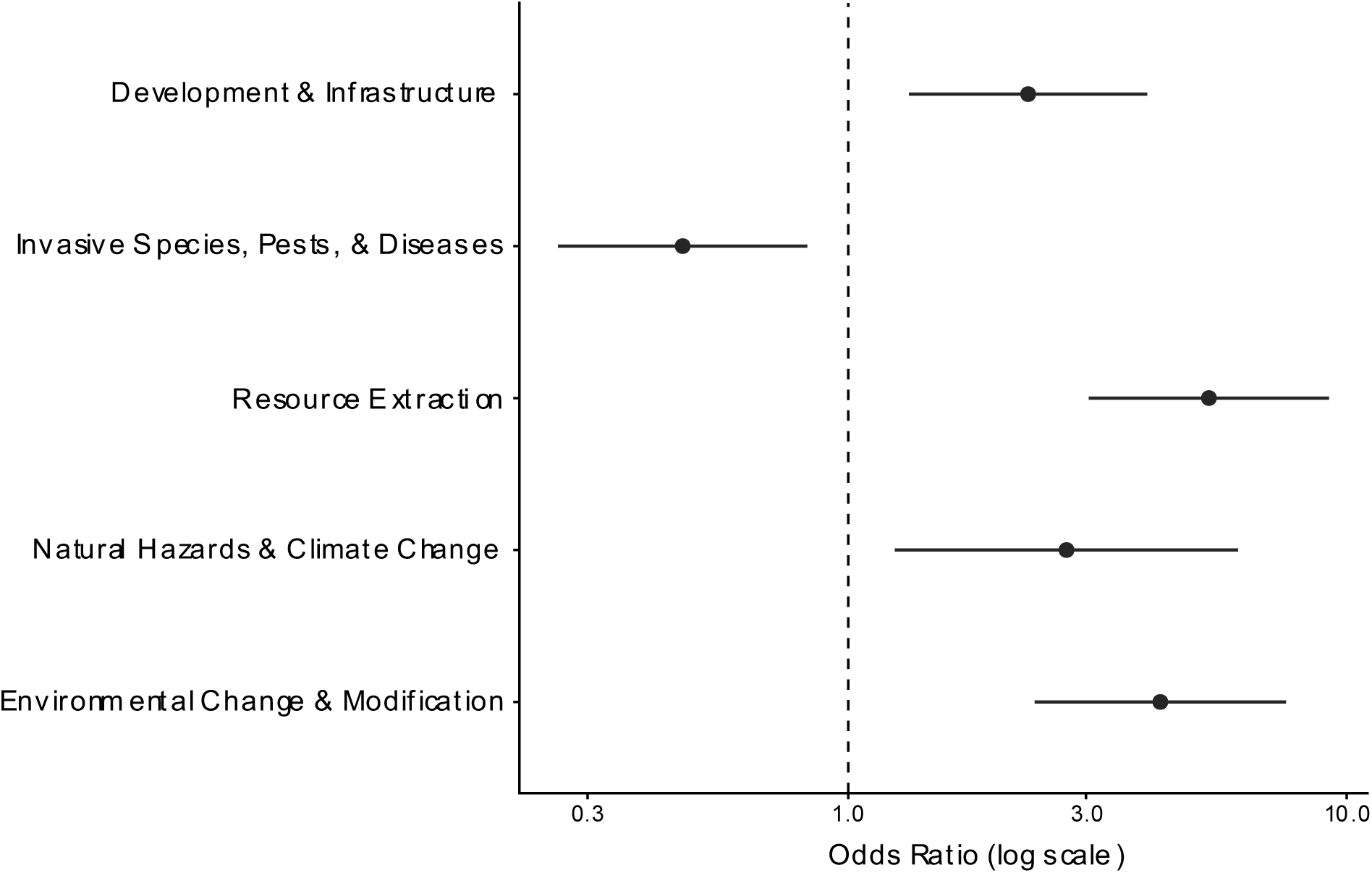
Forest plot of the odds ratios showing the effect of the presence of environmental pressures on the predicted probability of extinction risk.

**Table 7.**
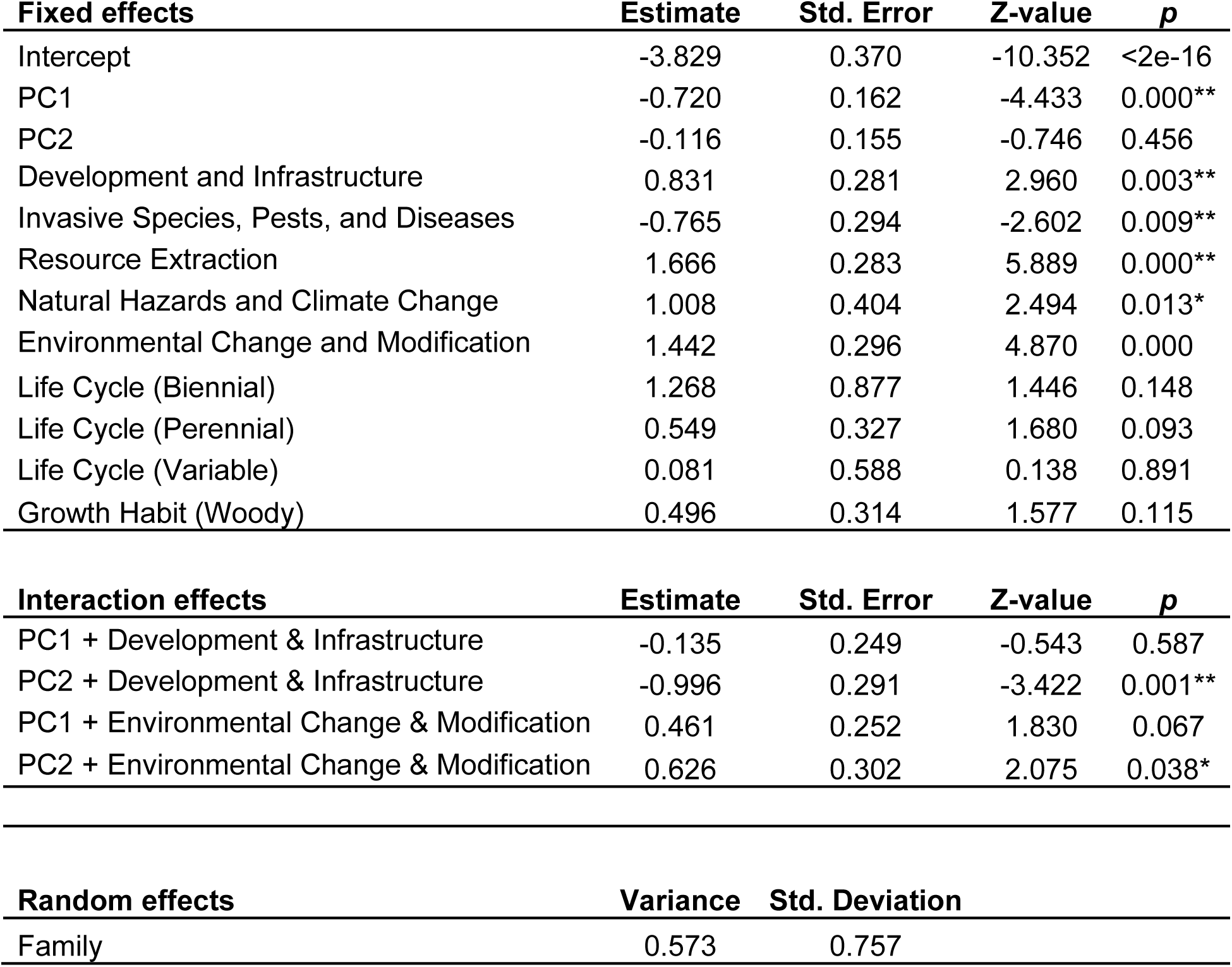
Summary results of generalised linear mixed-effect model predicting the effect of threats and their interaction with PC values on the probability of extinction risk.

| **Fixed effects** | **Estimate** | **Std. Error** | **Z-value** | **p** |
| --- | --- | --- | --- | --- |
| Intercept | -3.829 | 0.370 | -10.352 | <2e-16 |
| PC1 | -0.720 | 0.162 | -4.433 | 0.000** |
| PC2 | -0.116 | 0.155 | -0.746 | 0.456 |
| Development and Infrastructure | 0.831 | 0.281 | 2.960 | 0.003** |
| Invasive Species, Pests, and Diseases | -0.765 | 0.294 | -2.602 | 0.009** |
| Resource Extraction | 1.666 | 0.283 | 5.889 | 0.000** |
| Natural Hazards and Climate Change | 1.008 | 0.404 | 2.494 | 0.013* |
| Environmental Change and Modification | 1.442 | 0.296 | 4.870 | 0.000 |
| Life Cycle (Biennial) | 1.268 | 0.877 | 1.446 | 0.148 |
| Life Cycle (Perennial) | 0.549 | 0.327 | 1.680 | 0.093 |
| Life Cycle (Variable) | 0.081 | 0.588 | 0.138 | 0.891 |
| Growth Habit (Woody) | 0.496 | 0.314 | 1.577 | 0.115 |
| **Interaction effects** | **Estimate** | **Std. Error** | **Z-value** | **p** |
| PC1 + Development & Infrastructure | -0.135 | 0.249 | -0.543 | 0.587 |
| PC2 + Development & Infrastructure | -0.996 | 0.291 | -3.422 | 0.001** |
| PC1 + Environmental Change & Modification | 0.461 | 0.252 | 1.830 | 0.067 |
| PC2 + Environmental Change & Modification | 0.626 | 0.302 | 2.075 | 0.038* |
| **Random effects** | **Variance** | **Std. Deviation** |  |  |
| Family | 0.573 | 0.757 |  |  |

The effects of ‘*Development and Infrastructure’* and ‘*Environmental Change and Modification*’ are significantly modified by PC2, which represents leaf and fruit length. ‘*Development and Infrastructure’* significantly increased extinction risk for species with low PC2 values (short leaves, long fruits). Predicted probability of extinction risk increases from 5% to 30% as PC2 drops from 0 to -2, compared to a 1% increase in the absence of this threat (Figure 5b). ‘*Environmental Change and Modification*’ significantly increased extinction risk for species with high PC2 values (long leaves, short fruits), with the predicted probability increasing from 9% to 22% as PC2 values increase from 0 to 2; in the absence of this threat, there is no change (Figure 5d). We did not find significant trait and threat interactions for PC1 (Figure 5a, c).

**Figure 5.**
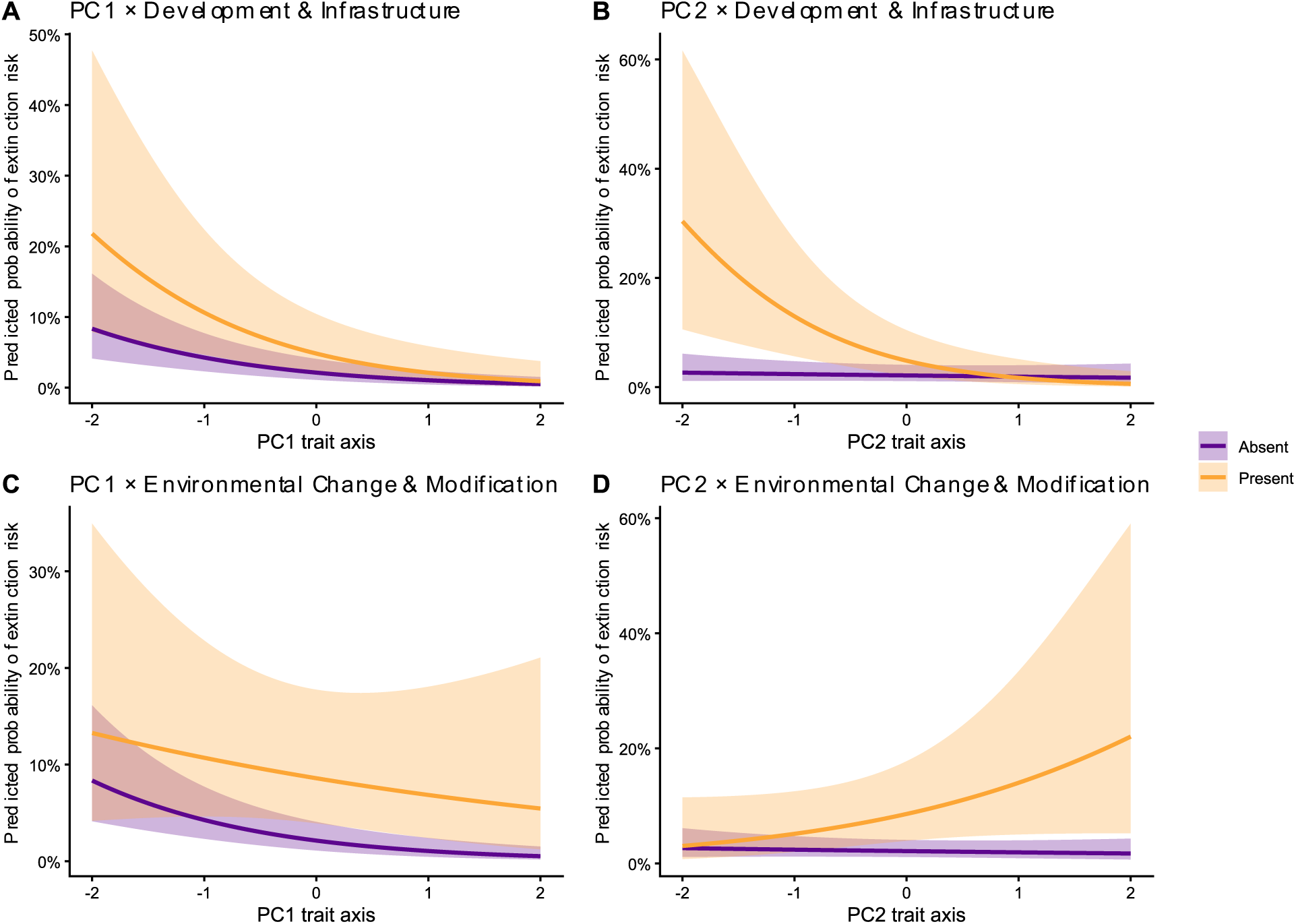
Predicted values of the interaction between the presence of threats and PC values on the probability of extinction risk.

## Discussion

Our understanding of how plant traits and human-induced threats interact to affect plant extinction risk is based on large-scale global analyses which use a limited number of traits (e.g. growth form, Bachman et al., 2024, Pelletier et al., 2018) or threats (e.g. climate change, Butt and Gallagher, 2018, Dudley et al., 2021; overhunting, Caughlin et al., 2015). While a critical starting point, such studies assume that trait-threat relationships using limited data averaged across broad spatial scales can capture regional processes where conservation efforts and outcomes are often focused. As a result, this approach limits our ability to use these trait-threat-risk frameworks to identify highly threatened species and prioritise conservation efforts. Using a broad range of plant functional traits for the angiosperm flora of Florida and a well-defined set of threats, we find that smaller-statured and herbaceous species (e.g. lower values on PC1) are at increased risk of extinction compared to larger, woody species (e.g. higher values on PC1). These smaller herbaceous species possess traits consistent with faster life-history strategies including rapid growth, high reproductive output, and higher dispersal capacity (Reich, 2014; Salguero-Gómez et al., 2016; Beckman et al., 2018). The effects of some human-induced threats are also modified by leaf and fruit length, but in opposing directions. Because a single trait can increase risk under one threat and lower it under another, plant traits therefore do not have fixed relationships to risk (Böhm et al., 2016). These results highlight the importance of incorporating a range of human-induced threats and functional traits, whilst also considering regional differences in floristic diversity, histories, and disturbance regimes for trait-based prediction to be a reliable conservation resource. Below, we explore how our results can improve trait-threat-risk frameworks and clarify how natural resilience and vulnerability vary under human-induced threats.

Within Florida, smaller, herbaceous species have an increased probability of extinction compared to larger, woody species. This result is the reverse of patterns found by previous studies, which uncovered that larger long-lived plants with slow life-history strategies are at greater risk of extinction due to increased generation times, low population turnover, and consequently reduced adaptive capacity to environmental changes and perturbations (Moles et al., 2009; Reich, 2014; Salguero-Gómez et al., 2016). We also find that species with long leaves and short fruits (high values on PC2), predominantly herbaceous species, are at a greater risk of extinction than other species under ‘*Environmental Change and Modification’*. This threat category encompasses human land management that leads to fire suppression and hydrological change such as water impoundment, both of which commonly occur across Florida (Abrahamson and Abrahamson, 1996; Abrahamson et al., 2021). These processes are likely to have a greater impact on plant groups such as Poaceae and Asteraceae, which for many native Florida species possess fire-adaptive traits and fire-induced reproductive triggers in subtropical and wet prairie grasslands (Main and Barry, 2002; Orzell et al., 2024). Fruit size has been found to strongly positively correlate with seed size and can serve as a proxy for seed size, therefore providing insights into dispersal mechanisms, dispersal capabilities (Stevenson et al., 2023), and seedling vigour and success (Baskin and Baskin, 2014). Species with smaller seeds are common within open and disturbed environments, including fire-maintained habitats, and require high levels of light availability to enable rapid growth and germination due to reduced nutritional seed reserves (Wendt et al., 2022).

Over time, fire suppression and hydrology modifications initiate shifts from open habitats towards more closed hardwood-dominated habitats, subsequently reducing the extent and persistence of native herbaceous taxa (Zambrano et al., 2019; Wieczorkowski and Lehmann, 2022; Matsuo et al., 2023; Smith et al., 2026). Within Florida’s longleaf pine woodlands and sand scrub habitats (NatureServe, 2026b,c), regional herbaceous endemic plant species (Kautz and Cox, 2001) such as *Pityopsis flexuosa* Small and habitat-dependent vertebrates (e.g. *Aphelocoma coerulescens Bosc*; NatureServe, 2026d) are threatened by the long-term consequences of fire suppression, with our results highlighting the ecosystem-level effects of these human-induced processes.

These results also reiterate the importance of habitat-specific fire prescription in natural areas of Florida, not only to reduce risk of uncontrollable wildfires but also as a conservation management practise to proxy natural ecosystem processes (Jones and Koptur, 2017; Connell et al., 2019; Morgan et al., 2020).

In the presence of ‘*Development and Infrastructure’*, plants with short leaves and long fruits (low values on PC2), predominantly woody shrubs and trees, have a higher extinction risk, the opposite of the relationship recovered in the presence of ‘*Environmental Change and Modification’*. Unlike ‘*Environmental Change and Modification’,* which incorporates threats generated by human activities such as hydrological and fire regime changes, which modify, disturb, or destroy habitats, the ‘*Development and Infrastructure’* threat incorporates the impacts of urban and transport land use changes. These processes often result in habitat loss and subsequently significant habitat fragmentation. Large-seeded species (long fruits) typically have longer generation times than other species and are more often dispersed by large-bodied mammals (Osuri and Sankaran, 2016; Beckman et al., 2018; Wendt et al., 2022). Habitat fragmentation can reduce mammal abundance and diversity in natural habitats by restricting or altering natural movement patterns, whilst dispersal of smaller-seeded species via wind or bird dispersal has been shown to increase in abundance (Cramer et al., 2007; Liu et al., 2019). As a result, habitat loss and fragmentation following land clearance disproportionately impact species with longer generation times and those which rely on animal vectors for dispersal, in agreement with patterns found by Moles et al. (2009), Reich (2014), and Salguero-Gómez et al. (2016) but in conflict with the relationship we recovered between traits and the effect of the threat ‘*Environmental Change and Modification’* on extinction risk.

Collectively, these results demonstrate differing responses of large-and small-seeded species to habitat loss and fragmentation. The varied interactions recovered here between threats and the same traits when considering ‘*Environmental Change and Modification’* and ‘*Development and Infrastructure’* are due to different impacts of these threats on habitat structure and processes, together with species-level niche differences. As a result, different growth habits and forms are disproportionately affected by different threats. Plant functional trait relationships can therefore be used to infer species-specific responses to different human-induced threats and provide insights for conservation managers. In model one, when only accounting for extinction risk and not threat type, we also find perennial species have an increased probability of extinction. This indicates that whilst life cycle contributes to overall species extinction risk, annuals and perennials are not comparatively disproportionately impacted by specific human-induced threats.

Apart from ‘*Invasive Species, Pests, and Diseases*’ (OR = 0.47), all threats were found to significantly increase extinction risk. ‘*Resource Extraction’*, which includes activities such as agriculture, livestock farming, and logging and wood harvesting, had the strongest effect on extinction risk (OR = 5.29). This reflects the economic importance and extent of agriculture and livestock farming (Swain et al., 2013; Campbell et al., 2022) and forestry industries (Martin et al., 2017) across the state. A similar increase in extinction risk is seen in the presence of ‘*Environmental Change and Modification’* threats (OR = 4.23). Both of these threat types enable, aid, and improve the expansion and subsistence of human populations, often resulting in habitat alteration, degradation, or loss (Sato and Lindenmayer, 2017). Therefore, with Florida’s human population projected to continue to rise (Rayer, 2026), the effects of threats will compound, putting the native flora of Florida and its ecosystems under increasing pressure (Raven and Wackernagel, 2020).

The one threat associated with lower extinction risk is ‘*Invasive Species, Pests, and Diseases*’ (OR = 0.47). We interpret this result as a likely positive impact of the management of invasive species such as *Dolichandra unguis-cati* (L.) L.G.Lohmann (cat’s-claw vine), *Eichhornia crassipes* (Mart.) Solms. (water hyacinth), and *Hydrilla verticillata* (L.f.) Royle (water thyme) in some areas of Florida, rather than a protective effect of this threat. The state of Florida has made substantial financial investment in managing invasive species to reduce their extent and impact, with funding targeted based on the prevalence of invasives and the conservation value of the habitat (Hiatt et al., 2019). For example, the extent of uncontrolled invasive plant species occurring in conservation areas within the state was reduced by a third from 1.5 million acres between 1995 and 2007 (Lee et al., 2009). Similarly, extensive research and management have reduced the extent of *Melaleuca quinquenervia* (Cav.) S.T.Blake from 400,000 ha to less than 100,000 ha across southern Florida including the Everglades, restoring natural ecological and environmental processes and subsequently preserving native biodiversity (Smith, 2022). In instances where management practices are directly addressing threats, the effect on extinction risk may reflect the response rather than the threat itself. It also highlights that the impact of human-induced disturbance and habitat loss can be mitigated, if not prevented, with informed conservation management and urban planning that can support economic sustainability whilst maintaining biodiversity.

By incorporating information on plant functional traits, which proxy key life-history and dispersal strategies, along with knowledge of regional ecosystem processes and human-induced threats, we can predict species-specific risk of extinction. Understanding these relationships will be beneficial when constructing conservation management plans and priorities for range-restricted and insufficiently studied taxa, which prevent a conservation assessment (Trethowan et al., 2025) with functional plant traits enabling predictions of extinction status. We expect regional and ecosystem differences in the relationships recovered among these plant functional traits, threats, and extinction risk, reflecting differing responses based on environmental effect, life histories, and specialised adaptations. As Florida is part of the larger North American Coastal Plain biodiversity hotspot (Noss et al., 2015), we do, however, expect our results to extrapolate across this region, given shared floristic compositions and habitats.

Despite extensive collaborative efforts and the establishment of downloadable trait databases (e.g. TRY, BIEN, and GIFT), large global-scale data gaps still exist for many species (particularly rare and at-risk species) for key functional traits, including reproductive, phenological, and hydrological traits (Dudley et al., 2019; Carmona and Beccari, 2025; Grenié et al., 2025). Moreover, as highlighted here, available trait databases exhibit clear taxonomic biases (e.g. gymnosperms vs. angiosperms). Such data deficiencies will constrain any attempt to conduct analyses of trait-threat interactions at larger spatial extents. Strategically filling these gaps for species of conservation concern and for underrepresented regions and taxonomic groups will be one of the most direct ways to further extend the use of trait-threat-risk frameworks outlined here.

## Acknowledgements

This work was supported by the US National Science Foundation, grant number DEB-2325836 to PS, RG, and DS. The authors are grateful to Geoff Levin of the Flora of North America Association for sharing provisional and unpublished floristic accounts. We would also like to thank Julie Allen for her suggestions and comments on earlier versions of the manuscript, along with JT Miller for providing a species list of native seed plants in Florida.

## Author contributions

ZDC, DS, PS, and RPG conceived the study. ZDC and RLF assembled trait data. ZDC and RPG analysed the data. ZDC wrote the first version of the manuscript. All authors read, edited, and approved the final manuscript.

## Supporting Information

Additional supporting information can be found online at the end of this article.

## Data availability

Compiled trait and threat data, along with scripts used to conduct the analysis, are deposited at:

