## Supplementary Figures and Tables for "Regional trait-threat interactions determine extinction risk in plants"

**Supplementary Materials**

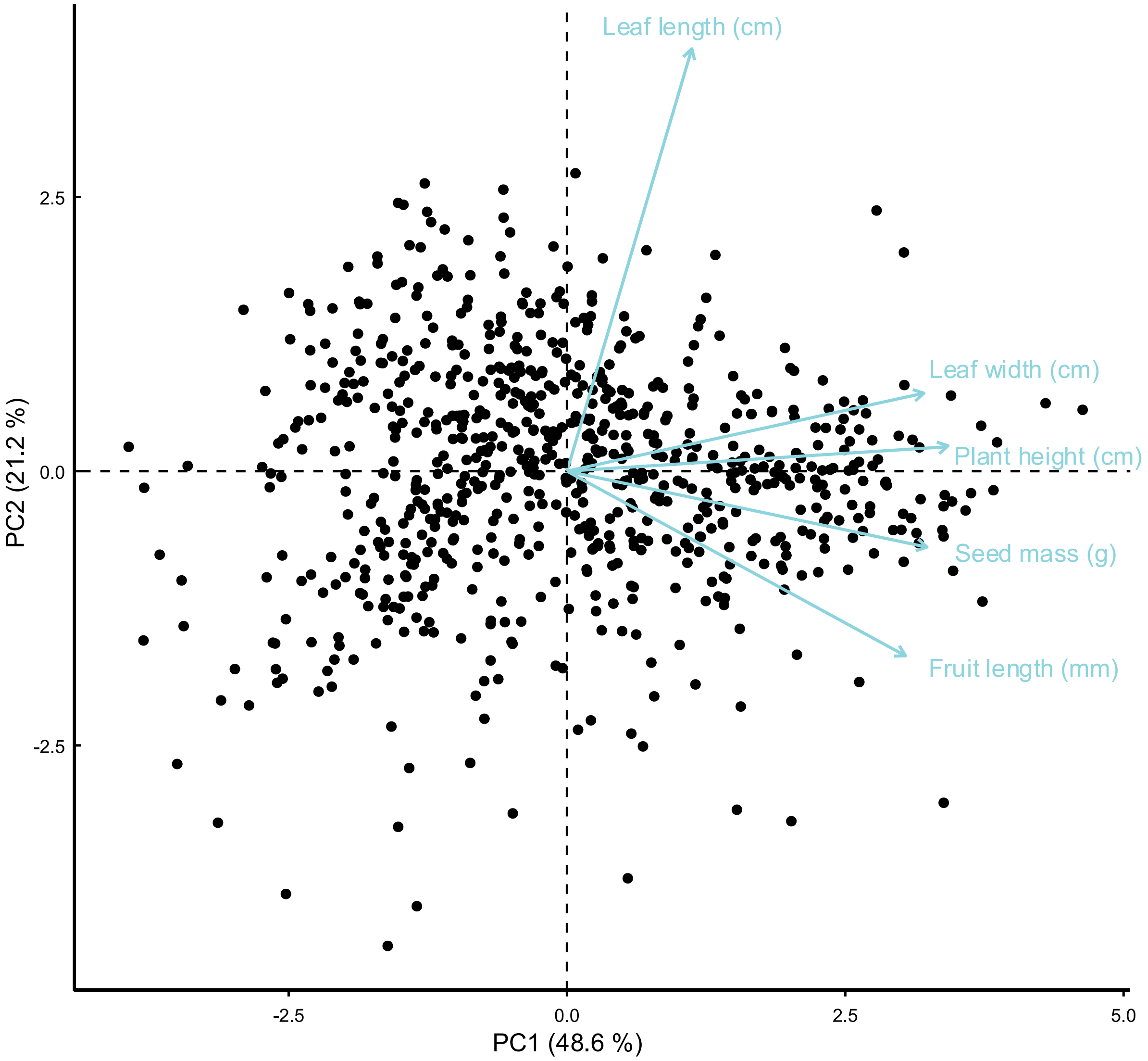


Supplementary Figure 1 – Principal component analysis biplot of trait variation for 691 plant species native to Florida with NatureServe assessments and available seed mass data. Continuous trait values are logged: fruit length (mm), leaf length (cm), leaf width (cm), plant height (cm), and seed mass (g). Each circle represents one taxon.

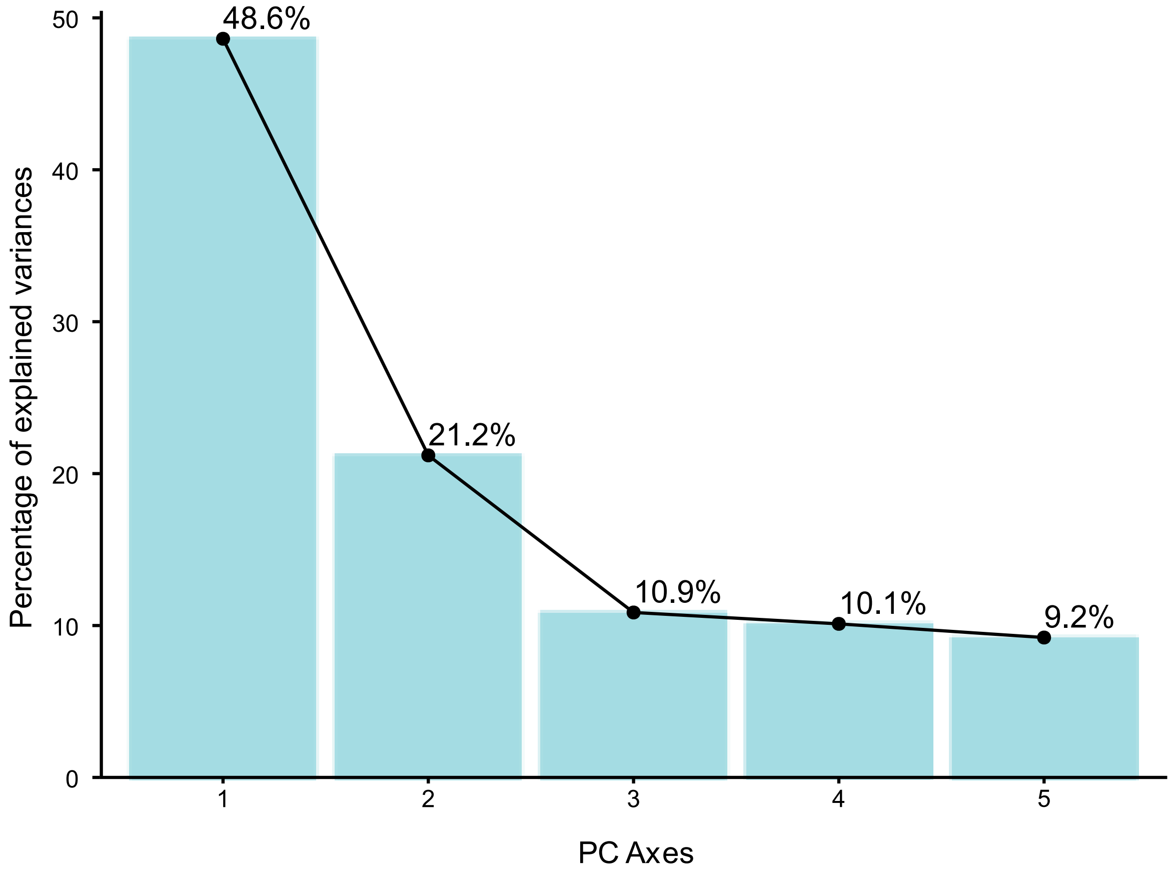


Supplementary Figure 2 – Principal component analysis eigenvalue scree plot of trait variation including seed mass data for 699 taxa native to Florida. Trait loadings for each PC axis are presented in supplementary table 5.

Supplementary Table 1 – Coverage of all plant trait data targeted as part of study, for all native plant taxa (percentages are shown in parentheses), along with source types (D = trait databases, F = Floras, O = Online resources).

| Measurement | Trait | Taxa represented | Sources |
|---|---|---|---|
| Categorical | **Dispersal Syndrome** | 729 (24.39%) | D |
| | **Fruit Type** | 2981 (99.73%) | F,D,O |
| | **Growth Form** | 2986 (99.90%) | D |
| | **Life Cycle** | 2987 (99.93%) | D |
| | **Growth Habit** | 2978 (99.63%) | D |
| Continuous | **Fruit Length (mm)** | 2288 (76.55%) | F,D,O |
| | **Fruit Width (mm)** | 1476 (49.38%) | F,D,O |
| | **Leaf Length (cm)** | 2499 (83.61%) | F,D,O |
| | **Leaf Nitrogen** | 540 (18.07%) | D |
| | **Leaf Thickness (cm)** | 366 (12.24%) | D |
| | **Leaf Width (cm)** | 2379 (79.59%) | D |
| | **Plant Height (cm)** | 2516 (84.18%) | F,D,O |
| | **Seed Length (mm)** | 927 (31.01%) | F,D,O |
| | **Seed Mass (mg)** | 1195 (39.98%) | F,D,O |
| | **Seed Width (mm)** | 631 (21.11%) | F,D,O |
| | **SLA (cmg)** | 659 (22.05%) | D |
| | **Wood Density (gcm3)** | 236 (7.90%) | D |

Supplementary Table 2 – List of sources, their type, and the method used to assemble trait data for all targeted traits. A short description for each source is provided.

| Source | Type | Method | Description | Link |
|---|---|---|---|---|
| **BIEN** | Database | R Package (bien) | Global database of plant functional traits and phylogenetic data, plot data, species occurrences, distributions, and checklists. | https://bien.nceas.ucsb.edu/bien/ |
| **Flora of North America** | Flora | Rule-based parsing (HTML) and manual extraction (PDF) | Floristic account of all plants native and naturalised in North America | https://floranorthamerica.org/Main_Page |
| **Flora of the Southeastern United States** | Flora | Manual look-up | Taxon accounts for all native and non-native taxa known to occur in the Southeastern United States. | https://fsus.ncbg.unc.edu/ |
| **GIFT** | Database | R Package (gift) | Global database of plant checklists, distribution, and functional traits. | https://biogeomacro.github.io/GIFT/ |
| **Global Wood Density Database** | Database | Downloadable database | Database of major wood functional traits for non-herbaceous taxa. | https://zenodo.org/records/18262736 |
| **Lady Bird Johnson Wildflower Database** | Database | Manual look-up | Native plants database of plants in North America (24,247 species accounts). | https://www.wildflower.org/plants/ |
| **Missouri Botanical Garden Plant Finder** | Database | Manual look-up | Taxon accounts for 7,500 plants growing or previously grown in the Kemper Centre display gardens. | https://www.missouribotanicalgarden.org/PlantFinder/PlantFinderSearch.aspx |
| **North Carolina Plant Toolbox** | Database | Manual look-up | Taxon accounts for 4,692 plants known to grow in and around North Carolina. | https://plants.ces.ncsu.edu/ |
| **PLANTS Database** | Database | Manual look-up | Database with standardised information across all vascular plants, mosses, liverworts, hornworts, and lichens of the United States and its territories | https://plants.sc.egov.usda.gov/ |
| **Plants of the World Online** | Database | Manual look-up | Plant taxonomy and descriptions for all plant names accepted by WCVP globally. | https://powo.science.kew.org/ |
| **Seed Information Database** | Database | Manual look-up | Database of seed biological trait data primarily from seed collections held in RBG Kew's Millennium Seed Bank | https://ser-sid.org/ |

| | | | | |
|---|---|---|---|---|
| **TRY Plant Trait Database** | Database | Downloadable database | Global open-access and source plant functional trait database. | https://www.try-db.org/ |
| **Vascular Plants of North Carolina** | Flora | Manual look-up | Taxon accounts for all native and non-native taxa known to occur in North Carolina. | https://auth1.dpr.ncparks.gov/flora/plant_list.php |
| **World Flora Online** | Flora | Manual look-up | Open-access compilation of all plant names and descriptions globally. | https://www.worldfloraonline.org/ |

Supplementary Table 3 – List of the Flora of North America volumes and families where taxon accounts provided where taxon accounts are currently not available online.

| Family | Number of native taxa in Florida | Volume |
|---|---|---|
| Anacardiaceae | 9 | 13 |
| Apiaceae | 37 | 13 |
| Aquifoliaceae | 11 | 18 |
| Araliaceae | 6 | 13 |
| Boraginaceae | 21 | 15 |
| Burseraceae | 2 | 13 |
| Caprifoliaceae | 4 | 18 |
| Geraniaceae | 3 | 13 |
| Lentibulariaceae | 20 | 18 |
| Menyanthaceae | 4 | 18 |
| Rubiaceae | 45 | 18 |
| Rutaceae | 9 | 13 |
| Sapindaceae | 11 | 13 |
| Viburnaceae | 5 | 18 |

Supplementary Table 4 – NatureServe Global Conservation Status ranks and descriptions. For the purpose of this study, taxa assigned global statuses between G1-G3 are considered at risk, and taxa ranked G4 or G5 are considered secure. Full definitions can be found online at: https://explorer.natureserve.org/AboutTheData/DataTypes/ConservationStatusCategories.

| Rank | Description | Study Conservation status |
|---|---|---|
| G1 | Critically Imperiled | At risk |
| G2 | Imperiled | |
| G3 | Vulnerable | |
| G4 | Apparently Secure | Secure |
| G5 | Secure | |
| GX | Presumed Extinct | Extinct |
| GH | Possibly Extinct | |

Supplementary Table 5 – Threat classifications used in this study and their corresponding IUCN-CMP Level 1 categories v.2.3 and summarised descriptions (IUCN, 2012). Full descriptions can be found at: https://nc.iucnredlist.org/redlist/content/attachment_files/dec_2012_guidance_threats_classification_scheme.pdf. Accessed on 7th July 2026

| Threat | IUCN-CMP Level 1 Categories | IUCN Summary Description |
|---|---|---|
| **Development and Infrastructure** | 1. Residential & Commercial Development | Threats from human settlements and non-agricultural land uses (e.g. domestic, commercial, and tourism development). |
| | 4. Transportation & Service Corridors | Threats from transport corridors and the use of them (e.g. roads, railroads, utility and service lines). |
| **Environmental Change and Modification** | 6. Human Intrusions & Disturbance | Threats from activities that cause habitat alteration, destruction, or disturbance (e.g. recreation, war). |
| | 7. Natural System Modifications | Threats from activities that cause habitat change, conversion, or degradation by managing natural and semi-natural systems for human welfare (e.g. fire management, hydrological change). |
| | 9. Pollution | Threats from the introduction of exotic or excessive levels of materials (e.g. wastewater, effluents). |
| **Invasive Species, Pests, and Diseases** | 8. Invasive & Other Problematic Species, Genes & Diseases | Threats from the non-native diseases, non-native species, and genes which result in biodiversity and population level losses. |
| **Natural Hazards and Climate Change** | 10. Geological Events | Threats from stochastic geological events (e.g. earthquakes, landslides, volcanic eruptions). |
| | 11. Climate Change & Severe Weather | Threats from long-term human-induced climatic change and weather events (e.g. temperature extremes, storms). |
| **Resource Extraction** | 2. Agriculture & Aquaculture | Threats from agricultural expansion and intensification including aquaculture, crop and livestock farming, and silviculture. |
| | 3. Energy Production & Mining | Threats from the extraction and use of non-biological resources (e.g. mineral, oil and gas extraction, and renewable energy). |
| | 5. Biological Resource Use | Threats from the use of wild biological resources (e.g. wild plant harvesting). |

Supplementary Table 6 – The contributions and correlations of each continuous trait with each principal component for 691 plant taxa native to Florida, which have available NatureServe assessments and seed mass data.

| | PC1 | PC2 | PC3 | PC4 | PC5 |
|---|---|---|---|---|---|
| | | | **Contributions** | | |
| **Plant Height (cm)** | 27.157 | 0.264 | 2.156 | 2.195 | 68.228 |
| **Fruit Length (mm)** | 21.560 | 15.152 | 3.124 | 47.168 | 12.996 |
| **Leaf Length (cm)** | 2.933 | 79.315 | 2.610 | 9.080 | 6.062 |
| **Leaf Width (cm)** | 23.966 | 2.705 | 49.501 | 22.647 | 1.181 |
| **Seed Mass (g)** | 24.383 | 2.564 | 42.609 | 18.911 | 11.533 |
| | | | **Correlations** | | |
| **Plant Height (cm)** | 0.813 | 0.053 | 0.108 | 0.105 | -0.561 |
| **Fruit Length (mm)** | 0.724 | -0.401 | -0.130 | 0.488 | 0.245 |
| **Leaf Length (cm)** | 0.267 | 0.917 | 0.119 | 0.214 | 0.167 |
| **Leaf Width (cm)** | 0.763 | 0.169 | -0.518 | -0.338 | 0.074 |
| **Seed Mass (g)** | 0.770 | -0.165 | 0.481 | -0.309 | 0.230 |

Supplementary Table 7 – AIC and weighted AIC scores for all models considered in our generalised linear mixed-model model framework.

| Model | Description | df | AIC | Delta AIC | Weighted Delta AIC |
|---|---|---|---|---|---|
| 1 | Traits only. | 8 | 1068.40 | 231.26 | 5.99E-51 |
| 2 | Cumulative threat as a continuous predictor. | 9 | 882.61 | 45.46 | 1.32E-10 |
| 3 | Cumulative threat as a factor. | 13 | 854.39 | 17.24 | 1.77E-04 |
| 4 | Binary threat presence. | 9 | 854.24 | 17.10 | 1.91E-04 |
| 5 | Binary threat for each category individually. | 13 | 845.62 | 8.47 | 1.43E-02 |
| 6 | Extended model 5 with interaction terms with PC axes and threat categories. | 17 | 837.15 | 0.00 | 9.85E-01 |

Supplementary Table 8 – Summary results of all models considered in our generalised linear mixed-effect model framework.

| Model 1 | | | | |
|---|---|---|---|---|
| **Fixed effects** | **Estimate** | **Standard Error** | **Z-value** | ***p*** |
| Intercept | -3.02671 | 0.30882 | -9.801 | < 2e-16*** |
| PC1 | -0.61156 | 0.11786 | -5.189 | 2.12e-07*** |
| PC2 | -0.24331 | 0.11309 | -2.151 | 0.03144* |
| Life Cycle (biennial) | 1.57634 | 0.76292 | 2.066 | 0.03881* |
| Life Cycle (perennial) | 0.85802 | 0.28294 | 3.033 | 0.00243** |
| Life Cycle (variable) | 0.01114 | 0.5211 | 0.021 | 0.98295 |
| Growth Habit (woody) | 0.2225 | 0.26858 | 0.828 | 0.40744 |
| **Random effects** | **Variance** | **Standard Deviation** | | |
| Family | 0.5357 | 0.7319 | | |

| Model 2 | | | | |
|---|---|---|---|---|
| **Fixed effects** | **Estimate** | **Standard Error** | **Z-value** | ***p*** |
| Intercept | -3.41026 | 0.35695 | -9.554 | < 2e-16*** |
| PC1 | -0.65314 | 0.12958 | -5.041 | 4.64e-07*** |
| PC2 | -0.2689 | 0.12933 | -2.079 | 0.0376* |
| Threat (cumulative) | 1.0073 | 0.08006 | 12.582 | < 2e-16*** |
| Life Cycle (biennial) | 1.30021 | 0.84151 | 1.545 | 0.1223 |
| Life Cycle (perennial) | 0.74908 | 0.32111 | 2.333 | 0.0197* |
| Life Cycle (variable) | 0.03327 | 0.57621 | 0.058 | 0.954 |
| Growth Habit (woody) | 0.36427 | 0.30047 | 1.212 | 0.2254 |
| **Random effects** | **Variance** | **Standard Deviation** | | |
| Family | 0.7087 | 0.8419 | | |

| Model 3 | | | | |
|---|---|---|---|---|
| **Fixed effects** | **Estimate** | **Standard Error** | **Z-value** | ***p*** |
| Intercept | -4.05369 | 0.36766 | -11.026 | < 2e-16*** |
| PC1 | -0.62641 | 0.12714 | -4.927 | 8.35e-07*** |
| PC2 | -0.24933 | 0.12625 | -1.975 | 0.0483* |
| Threat 1 (cumulative factor) | 2.34513 | 0.31433 | 7.461 | 8.61e-14*** |
| Threat 2 (cumulative factor) | 2.51264 | 0.26127 | 9.617 | < 2e-16*** |
| Threat 3 (cumulative factor) | 2.80553 | 0.27672 | 10.139 | < 2e-16*** |
| Threat 4 (cumulative factor) | 2.72633 | 0.35975 | 7.578 | 3.50e-14*** |
| Threat 5 (cumulative factor) | 4.1714 | 0.69414 | 6.009 | 1.86e-09*** |
| Life Cycle (biennial) | 1.16594 | 0.85504 | 1.364 | 0.1727 |
| Life Cycle (perennial) | 0.61989 | 0.32036 | 1.935 | 0.053 |
| Life Cycle (variable) | -0.02781 | 0.58965 | -0.047 | 0.9624 |
| Growth Habit (woody) | 0.35405 | 0.29898 | 1.184 | 0.2363 |
| **Random effects** | **Variance** | **Standard Deviation** | | |
| Family | 0.4894 | 0.6996 | | |

| Model 4 | | | | |
|---|---|---|---|---|
| **Fixed effects** | **Estimate** | **Standard Error** | **Z-value** | ***p*** |
| Intercept | -4.04634 | 0.35785 | -11.307 | < 2e-16*** |
| PC1 | -0.6294 | 0.12458 | -5.052 | 4.36e-07*** |
| PC2 | -0.2422 | 0.1238 | -1.956 | 0.0504 |
| All Threats (binarised) | 2.64662 | 0.1969 | 13.441 | < 2e-16*** |
| Life Cycle (biennial) | 1.16694 | 0.85138 | 1.371 | 0.1705 |
| Life Cycle (perennial) | 0.61401 | 0.31363 | 1.958 | 0.0503 |
| Life Cycle (variable) | 0.03924 | 0.57595 | 0.068 | 0.9457 |
| Growth Habit (woody) | 0.39526 | 0.29172 | 1.355 | 0.1754 |
| **Random effects** | **Variance** | **Standard Deviation** | | |
| Family | 0.4478 | 0.6692 | | |

| Model 5 | | | | |
|---|---|---|---|---|
| **Fixed effects** | **Estimate** | **Standard Error** | **Z-value** | ***p*** |
| Intercept | -3.7626 | 0.3603 | -10.443 | < 2e-16*** |
| PC1 | -0.5995 | 0.1302 | -4.605 | 4.13e-06*** |
| PC2 | -0.2692 | 0.1283 | -2.097 | 0.036* |
| Development and Infrastructure | 1.0355 | 0.2659 | 3.895 | 9.84e-05*** |
| Invasive Species, Pests, and Diseases | -0.7345 | 0.2883 | -2.547 | 0.0109* |
| Resource Extraction | 1.6707 | 0.2706 | 6.173 | 6.69e-10*** |
| Natural Hazards and Climate Change | 0.9453 | 0.3958 | 2.388 | 0.0169* |
| Environmental Change and Modification | 1.1831 | 0.2805 | 4.218 | 2.46e-05*** |
| Life Cycle (biennial) | 1.3069 | 0.8744 | 1.495 | 0.135 |
| Life Cycle (perennial) | 0.5442 | 0.3238 | 1.681 | 0.0928 |
| Life Cycle (variable) | 0.1217 | 0.5851 | 0.208 | 0.8352 |
| Growth Habit (woody) | 0.4453 | 0.304 | 1.465 | 0.1429 |
| **Random effects** | **Variance** | **Standard Deviation** | | |
| Family | 0.5285 | 0.727 | | |

| Model 6 | | | | |
|---|---|---|---|---|
| **Fixed effects** | **Estimate** | **Standard Error** | **Z-value** | **p** |
| Intercept | -3.829 | 0.370 | -10.352 | <2e-16*** |
| PC1 | -0.720 | 0.162 | -4.433 | 9.28e-06*** |
| PC2 | -0.116 | 0.155 | -0.746 | 0.4555 |
| Development and Infrastructure | 0.831 | 0.281 | 2.960 | 0.003077** |
| Invasive Species, Pests, and Diseases | -0.765 | 0.294 | -2.602 | 0.009272** |
| Resource Extraction | 1.666 | 0.283 | 5.889 | 3.89e-09*** |
| Natural Hazards and Climate Change | 1.008 | 0.404 | 2.494 | 0.012637* |
| Environmental Change and Modification | 1.442 | 0.296 | 4.870 | 1.12e-06*** |
| Life Cycle (biennial) | 1.268 | 0.877 | 1.446 | 0.1481 |
| Life Cycle (perennial) | 0.549 | 0.327 | 1.680 | 0.0929 |
| Life Cycle (variable) | 0.081 | 0.588 | 0.138 | 0.8906 |
| Growth Habit (woody) | 0.496 | 0.314 | 1.577 | 0.1148 |
| **Interaction effects** | **Estimate** | **Standard Error** | **Z-value** | **p** |
| PC1 + Development and Infrastructure | -0.135 | 0.249 | -0.543 | 0.58692 |
| PC2 + Development and Infrastructure | -0.996 | 0.291 | -3.422 | 0.000621*** |
| PC1 + Environmental Change and Modification | 0.461 | 0.252 | 1.830 | 0.06731 |
| PC2 + Environmental Change and Modification | 0.626 | 0.302 | 2.075 | 0.037947* |
| **Random effects** | **Variance** | **Standard Deviation** | | |
| Family | 0.573 | 0.757 | | |

Supplementary Table 9 – Variance inflation factors (VIF) for model one and model six generalised linear mixed effect models.

Model 1

| Fixed effects | GVIF | Df | GVIF^(1/(2*Df)) |
|---|---|---|---|
| PC1 | 1.4851 | 1 | 1.218647 |
| PC2 | 1.16638 | 1 | 1.079989 |
| Life Cycle | 1.17298 | 3 | 1.026949 |
| Growth Habit | 1.30743 | 1 | 1.143429 |

Model 6

| Fixed effects | GVIF | Df | GVIF^(1/(2*Df)) |
|---|---|---|---|
| PC1 | 2.21628 | 1 | 1.488716 |
| PC2 | 1.56422 | 1 | 1.250688 |
| Development and Infrastructure | 1.88041 | 1 | 1.371279 |
| Invasive Species, Pests, and Diseases | 1.55178 | 1 | 1.245705 |
| Resource Extraction | 1.52154 | 1 | 1.233506 |
| Natural Hazards and Climate Change | 1.19063 | 1 | 1.091161 |
| Environmental Change and Modification | 1.90529 | 1 | 1.380321 |
| Life Cycle | 1.22124 | 3 | 1.033872 |
| Growth Habit | 1.33608 | 1 | 1.155888 |
| **Interaction effects** | **GVIF** | **Df** | **GVIF^(1/(2*Df))** |
| PC1 + Development and Infrastructure | 2.09326 | 1 | 1.44681 |
| PC2 + Development and Infrastructure | 2.50873 | 1 | 1.583896 |
| PC1 + Environmental Change and Modification | 2.0423 | 1 | 1.429092 |
| PC2 + Environmental Change and Modification | 2.38116 | 1 | 1.5431 |
